# CBR4 is essential for mice but not for skeletal muscle function

**DOI:** 10.64898/2026.07.13.738135

**Authors:** Ali J. Masud, Guangyu Jiang, Kaija J. Autio, M. Tanvir Rahman, Irene M.G.M. Hemel, J. Kalervo Hiltunen, Alexander J. Kastaniotis

## Abstract

The mitochondrial fatty acid synthesis (mtFAS) pathway is a highly conserved process in mitochondria implicated in metabolic state sensing. Aberrant functioning of this pathway leads to neurodegenerative diseases in humans. Animal experiments indicates that the mtFAS pathway is essential in mammals, and mtFAS inactivation leads to neuronal cell death. Nuclear encoded mitochondrial 3-ketoacyl-acyl carrier protein reductase (KAR) is a heterotetrameric enzyme in this process, consisting of two CBR4 and two HSD17B8 polypeptides. CBR4 works as the catalytic subunit of the enzyme. Here, we provide evidence that CBR4 function is essential in mammals. In contrast, a skeletal muscle-specific *Cbr4* KO in mice did not result in any measurable defects in muscle strength and endurance, and the overall structure of the muscle remained unchanged. The *Cbr4* KO did not affect the lipoylation process in quadriceps muscle samples, and high-resolution respirometry analysis of soleus muscle samples showed no defects in mitochondrial respiration capacity. The lack of a phenotype of a muscle-specific *Cbr4* KO is consistent with previous reports on a lack of effects of mtFAS inactivation in muscle and re-iterates the question about the existence of bypass mechanisms that can alleviate mtFAS deficiencies in non-neuronal cell types.

## Introduction

For a long time, it was accepted that the well-characterized fatty acid (FA) synthesis pathway of non-plant eukaryotes takes place in the cytosol. This dogma has been refuted by data collected during the past three decades, indicating that eukaryotic mitochondria are also able to synthesize FAs. This synthesis pathway in mitochondria is now termed the “mitochondrial fatty acid synthesis” (mtFAS) pathway. Accumulating evidence supports the hypothesis that the mtFAS pathway acts as a sensor of cellular metabolic state, coordinating mitochondrial function and biogenesis with acetyl-CoA availability (1,2). In humans, defects in the gene encoding mtFAS enzyme MECR (mitochondrial 2-enoyl-CoA/ACP reductase) leads to MEPAN (Mitochondrial Enoyl Reductase Protein Associated Neurodegeneration) (3). The mtFAS pathway contributes to mitochondrial functions through its acyl products, tethered to acyl-carrier-protein (acyl-ACP) via two different mechanisms: (i) Acyl-ACPs physically interacting with Lys-Tyr-Arg (LYR)-motif proteins to form complexes that are indispensable for iron-sulfur cluster biosynthesis, mitochondrial respiratory chain complex I function and integrity, and assembly of complexes II, III and V as well as mitoribosomes and (ii) Providing the mitochondrially synthesized octanoyl-ACP precursor used in the synthesis of lipoic acid, a cofactor of mitochondrial pyruvate dehydrogenase (PDH), α-ketoglutarate dehydrogenase (αKGDH), branched-chain α-keto acid dehydrogenase (BCKDH), and the glycine cleavage system (GCSH) complexes (4). In contrast to the cytosolic fatty acid synthesis (FAS) pathway, where a single multifunctional enzyme complex in mammals catalyzes all the reactions, the synthesis in mtFAS proceeds by discrete enzymes.

Seven different nuclear-encoded proteins are involved in the acyl chain elongation process, and three different enzymes are involved in the transfer of the acyl group to the target protein (4). The mitochondrial 3-ketoacyl-ACP reductase (Oar1/KAR) is one of the enzymes in this cyclic pathway, catalyzing the first reduction reaction of 3-ketoacyl-ACP to 3-hydroxyacyl-ACP in yeast/mammals (5). The mammalian KAR is a heterotetrameric enzyme composed of two subunits, 17β-hydroxysteroid dehydrogenase type-8 (HSD17B8, α-subunit) and carbonyl reductase type-4 (CBR4, β-subunit) (5) whereas the fungal Oar1 assembles into a homotetrameric enzyme (6). CBR4 is the catalytic subunit of KAR, while HSD17B8 serves as a scaffold protein required for CBR4 stability and solubility (7). Both protein subunits showed strong interaction *in vitro* and *in vivo*. Their co-expression in yeast can complement the respiratory deficient phenotype of the yeast Δ*oar1* strain (5) and the proteins co-purify in a stable complex when co-expressed in *E. coli* (7).

In an earlier study CBR4 had been characterized as a carbonyl reductase required to protect cells from 9,10-phenanthenequinone-induced cytotoxicity (8). Later results by Chen *et al*., however, indicated that the true physiological role of CBR4 in mammals is to serve as catalytic subunit of human mitochondrial KAR (5). Apart from these studies, there is not much information about the physiological role of CBR4 in mammalian mitochondrial function. In the work presented here, we describe the investigation of the consequences of systemic loss of CBR4 in a mouse model using “Knockout-first” type *Cbr4* mutant mice, as well as for muscle-specific loss of CBR4 in an inducible skeletal muscle-specific *Cbr4* knockout (KO) mouse line. Together our data suggests that while ubiquitous knock out of *Cbr4* is embryonically lethal, conditional *Cbr4* knock out is dispensable in a muscle tissue specific context in post-weaning mice.

## Results

### A complete loss of *Cbr4* is lethal in mouse

At the start of this work, mammalian KAR KO models were unavailable. To study the role of *Cbr4* in mice, our first aim was to generate a mouse model with complete inactivation of *Cbr4* in all tissues and cells by deleting the exon 2 of *Cbr4*. Breeding of 12 heterozygous *Cbr4*^tm1d^ mouse couples led to birth of 59 pups. Out of those 40 pups were heterozygous and 19 were wild type. However, we were unable to obtain any homozygous *Cbr4*^tm1d^ mice. This result is consistent with an essential function of *Cbr4* in embryonic development. It mirrors the data obtained for KOs of *Mecr* or *NdufabI*, encoding human ACP (9,10), which also lead to embryonic lethality. All this data confirms that intact mtFAS is essential for embryonic development.

### Skeletal muscle-specific *Cbr4* KO mice

Since complete loss of *Cbr4* led to embryonal lethality, we decided to generate an inducible “loss of *Cbr4*” model initiated post-weaning and restricted to one tissue. Skeletal muscle was selected as the target due to the need for high levels of mitochondrial function and ATP in muscle. *Cbr4*^tm1c^ mice were bred with a mouse line carrying a construct that allows for doxycycline (DOX)-inducible, skeletal muscle-specific expression of Cre recombinase in HSA-rtTA/TRE-Cre. This mouse line allowed for induction of *Cbr4* loss in skeletal muscle at the age of six weeks, initiated by DOX ingestion. As a means of delivery of the inducing compound, these mice received 2 mg/ ml DOX in their drinking water, sweetened with 5% sucrose for better acceptance, for three weeks. At the same time, the control group was supplied with drinking water only laced with 5% sucrose. After the three-week induction period, all mice received regular drinking water until the end of the experiment.

### *Cbr4* mRNA gene expression level in quadriceps muscle

First after euthanizing the mice, we assessed the effect of doxycycline treatment for *Cbr4* expression in quadriceps muscle. A previous study showed that HSA-rtTA/TRE-Cre transgene is specifically expressed in the skeletal muscle but not in the heart, bladder, liver, kidney, spleen, or spinal cord tissues (11). As in previously published reports, our *Cbr4* conditional, skeletal muscle-specific mouse model, administration of DOX was used to stimulate the deletion of exon 2 of the *Cbr4* genomic DNA, leading to a transcript lacking this exon. Splicing between exon 1 and exon 3 of the *Cbr4* gene leads to an out-of-frame fusion, resulting in the translation of several missense codons, followed by a nonsense mutation leading to premature translation termination and a severely truncated protein. We were unable to obtain antibodies against CBR4 with any specificity for the protein in mouse tissue or cells to confirm the loss of Cbr4 protein in skeletal muscles. Therefore, we designed droplet digital PCR (ddPCR) primers (Table 1) specific for exon 2 of *Cbr4*. The ddPCR analysis showed that the female DOX+ mice had a clear and consistent reduction of *Cbr4* mRNA compared to DOX– control littermates (Fig. 1). The same trend was detected also in male mice, but there the different in expression levels did not reach the statistically significant level (Fig. 1). Therefore, we only included mice that showed at least 75% decrease in expression level of *Cbr4* in muscle tissue in our analyses, which were restricted to four treated and five control females.

**Table 1.**
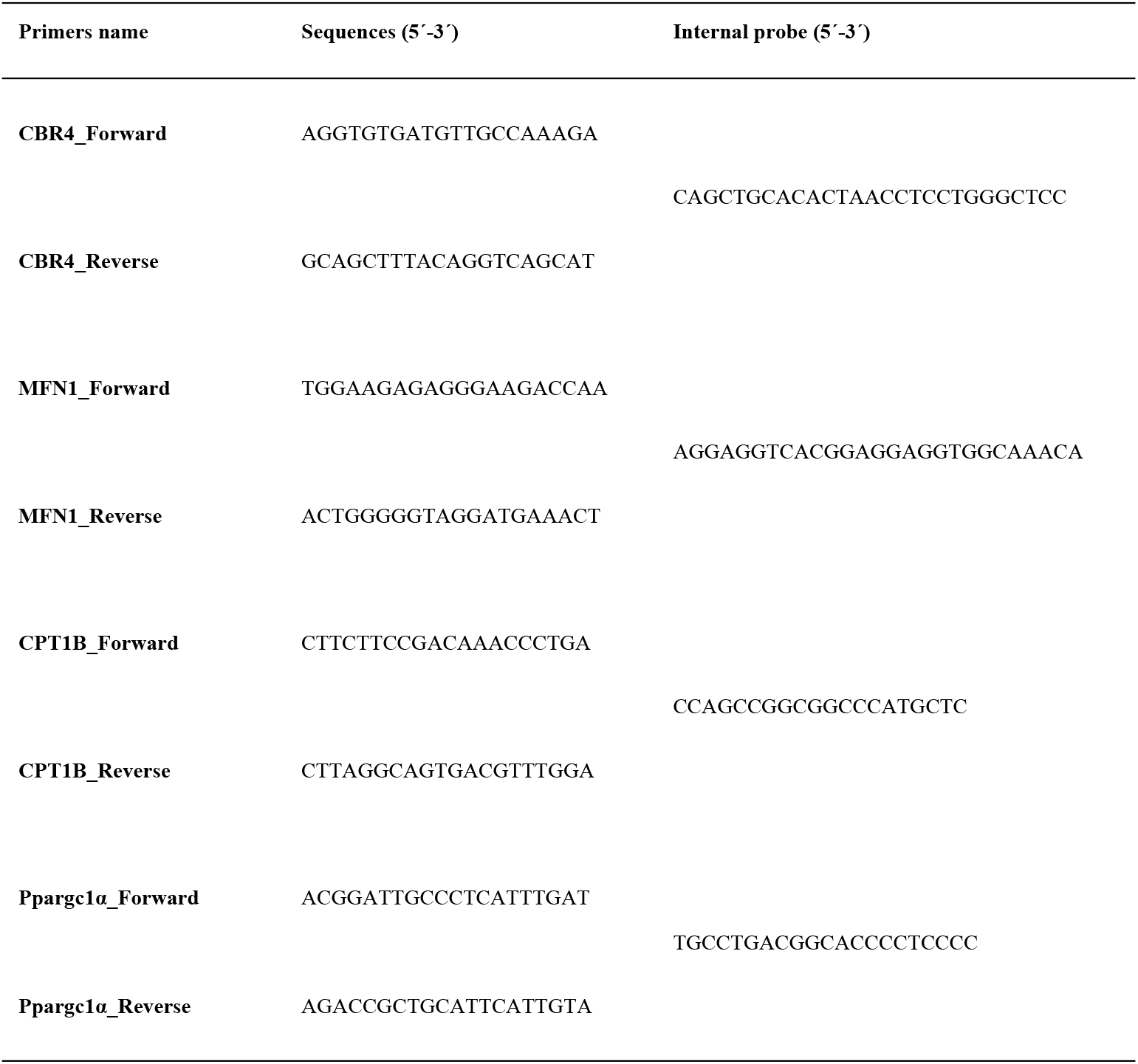
List of ddPCR primers used in the study.

**Fig. 1.**
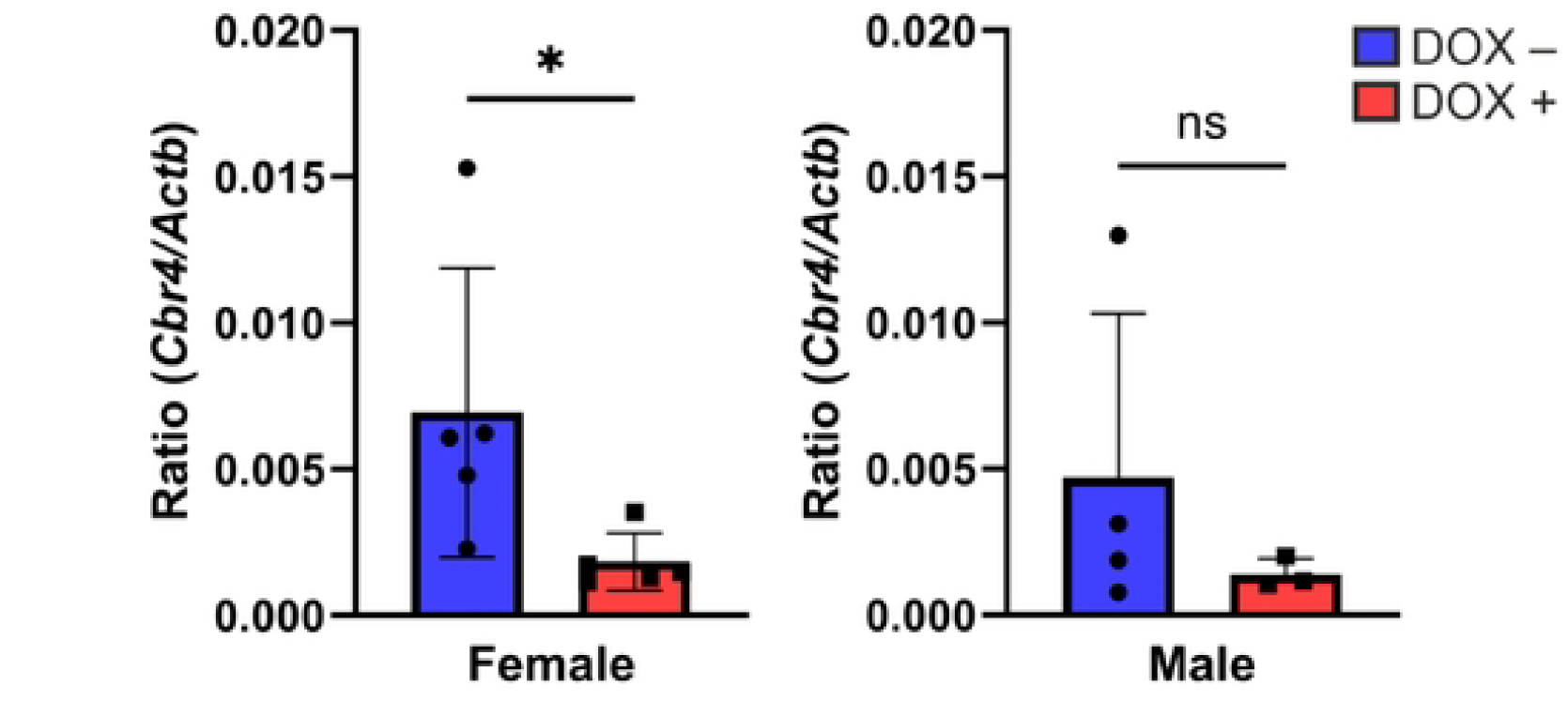
*Cbr4* mRNA expression in the quadriceps muscle of skeletal muscle. *Cbr4* mRNA expression in the quadriceps muscle of skeletal muscle-specific *Cbr4* KO mice was quantified using droplet digital PCR. *Cbr4* gene expression level was normalized against actin (*Actb*) and presented as *Cbr4/Actb* ratio. Doxycycline-administrated (DOX+) mice were compared to their sucrose-fed control littermates (DOX–) for females (n=5 per group) and males (n=4 DOX– and n=3 DOX+). A significant difference in Cbr4 expression was found for the female mice, however this was not the case for the male mice, where the *Cbr4* expression differed more in the DOX– controls. Groups were compared using a Wilcoxon signed-rank test, with p< 0.05 considered as significant. Data are presented as mean (±) SD, with *p<0.05 and ns=not significant.

No differences in weight gain in these mice were detected during the monitoring period for up to 48 weeks (Fig. 2A). The muscle performance was analyzed every second week employing a grid hanging test (Fig 2B). In addition, the maximal force the mice were able to exert in their fore limbs was tested at the age of 12 months by grip strength analysis (Fig 2C). Also, this approach did not reveal any differences between the animal groups. Two running tests to evaluate mice endurance were performed at the age of 11 months. In the first test all the mice were able to run 14 minutes with accelerating speed. The glucose and lactate levels in the blood were analyzed before, immediately after, and 30 minutes after running. Blood glucose levels between the mouse groups were indistinguishable independently of any condition tested (Fig 2D). Interestingly, DOX+ mice had elevated lactate levels before the running, but after the running exercise no differences in blood lactate were detected (Fig 2E). Finally, the running test until exhaustion was carried out, and here no differences in running performance were observed (Fig. 2F). The mice were euthanized at the age of 13 months.

**Fig 2.**
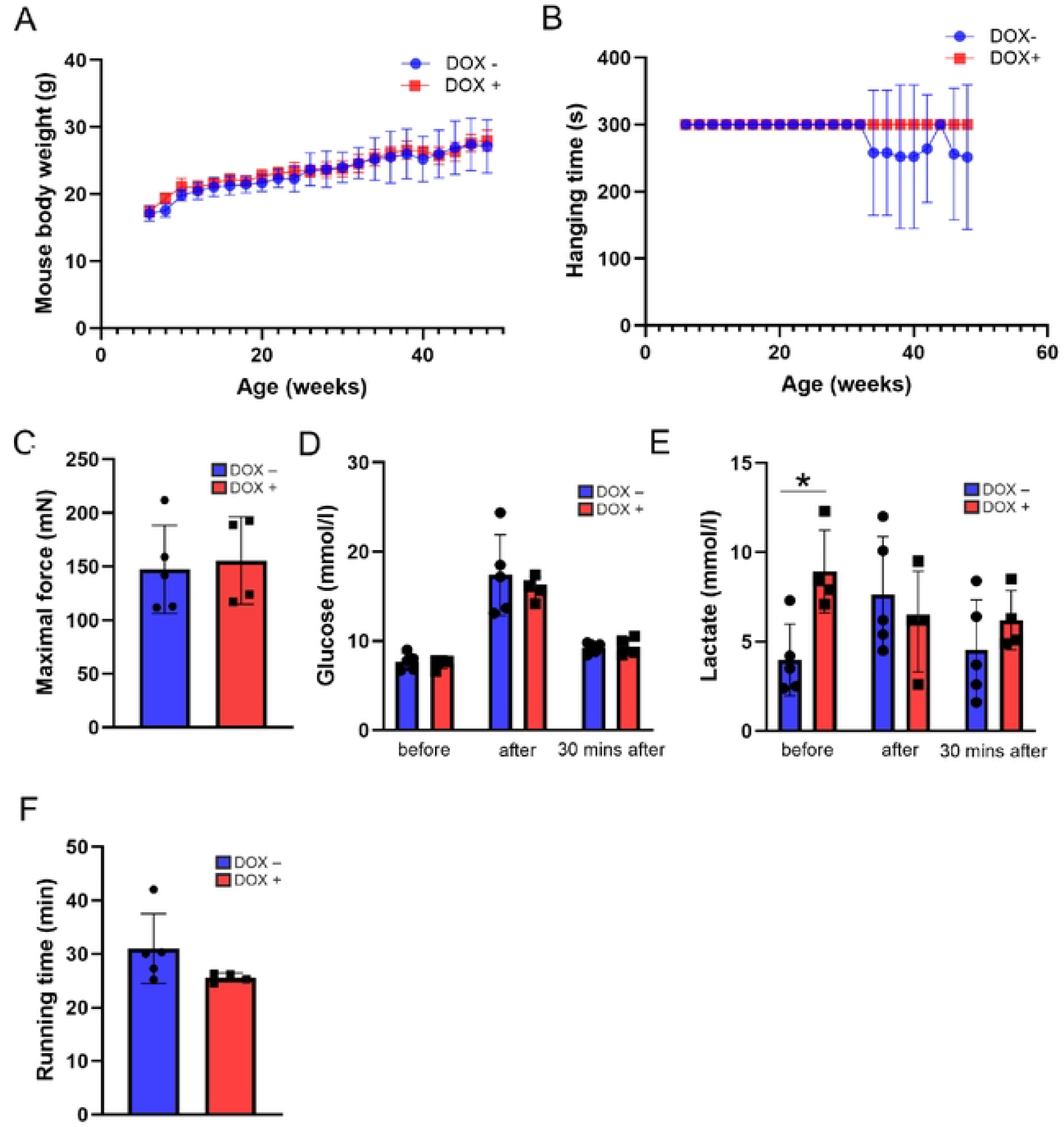
Effects of reduced *Cbr4* mRNA expression in skeletal muscle on female mouse performance. (A) Mouse body weights of sucrose fed (DOX–) and doxycycline induced (DOX+) littermate mice. Both DOX– and DOX+ mice continued to gain weight over the study period. (B) Grid hanging test of DOX– and DOX+ littermate mice where DOX– and DOX+ mice showed similar grid hanging abilities. All the mice were hanging for five minutes except one DOX– mouse that nearly always dropped after 1-2 minutes from week 34 onwards. (C) Maximal force of DOX– and DOX+ mice was analyzed with Grip Strength Meter and differences between the groups were detected. (D) Blood glucose and (E) lactate levels were measured before, immediately after and 30 minutes after 14 minutes running test in DOX– and DOX+ mice. The lactate concentration in blood was elevated in skeletal muscle-specific *Cbr4* KO mice before running, but in other conditions no changes were detected. Similarly, the glucose levels were indistinguishable in DOX– and DOX+ mice. (F) Both DOX– and DOX+ mice were capable to run about 30 minutes with increasing speed and no differences with the groups were noticed. The number of mice in experiments were DOX– (n=5) and DOX+ (n=4). Data are presented as mean (±) SD.

### Histology of the quadriceps muscle

The skeletal muscle morphology of the DOX– and DOX+ mice was analyzed from the H&E-stained quadriceps muscle sections. No visible defects were detected in the skeletal muscle morphology of the DOX+ mice when compared to the DOX– mice (Fig. 3A). Further quantification of the fiber cross sectional area (FCSA) of the quadriceps muscle showed no abnormality in the muscle morphology (Fig. 3B).

**Fig. 3.**
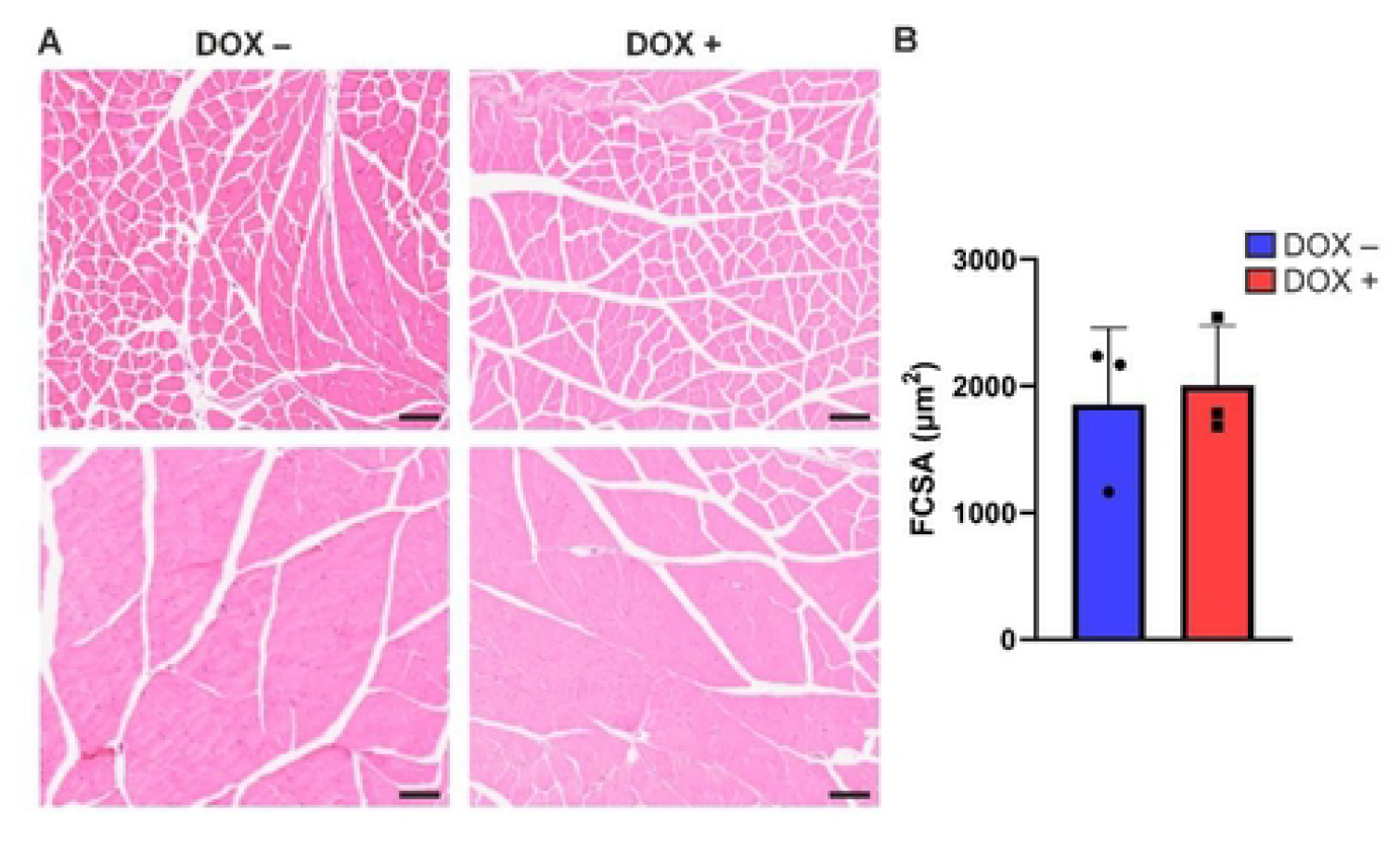
Morphological assessment of quadriceps muscle of skeletal muscle-specific Cbr4 KO mice. (A) H&E staining of quadriceps muscle sections of DOX– and DOX+ mice. Scale bars 100 μm. (B) Quantification of the fiber cross-sectional area (FCSA) of quadriceps muscle, n=3 mice per group. No difference in FCSA was found between DOX– and DOX+ mice. Groups were compared using a Wilcoxon signed-rank test, with p< 0.05 considered as significant. Data are presented as mean (±) SD.

### Mitochondrial respiratory chain activities in the soleus muscle

The effects of *Cbr4* KO on skeletal muscle mitochondrial respiratory complexes were analyzed using a substrate/inhibitor titration *ex vivo* approach in an OROBOROS Oxygraph-2K. Samples of soleus muscle, which consists mainly of type I muscle fibers with high respiratory activity, were used in the study. Substrates targeting the CI, CII and CIV linked OXPHOS were used to investigate their activities. Our substrate/inhibitor-based titration approach did not yield in significance differences between the DOX– and DOX+ mice (Fig. 4). Respiratory control ratio (RCR) is the ratio of respiration rate before and after the addition of a saturating concentration of ADP and indicates the coupling of respiration and OXPHOS. The RCR values for the DOX– and DOX+ samples were similar (Fig. 4F). The addition of cyt C did not show significant stimulation of respiration in DOX– or DOX+ mice indicating intactness of the OMM (Supplementary Fig. 1). On the other hand, addition of ADP caused high activation of respiration in both DOX– and DOX+ mice samples, indicating successful permeabilization of soleus muscle by saponin (Supplementary Fig. 1).

**Fig. 4.**
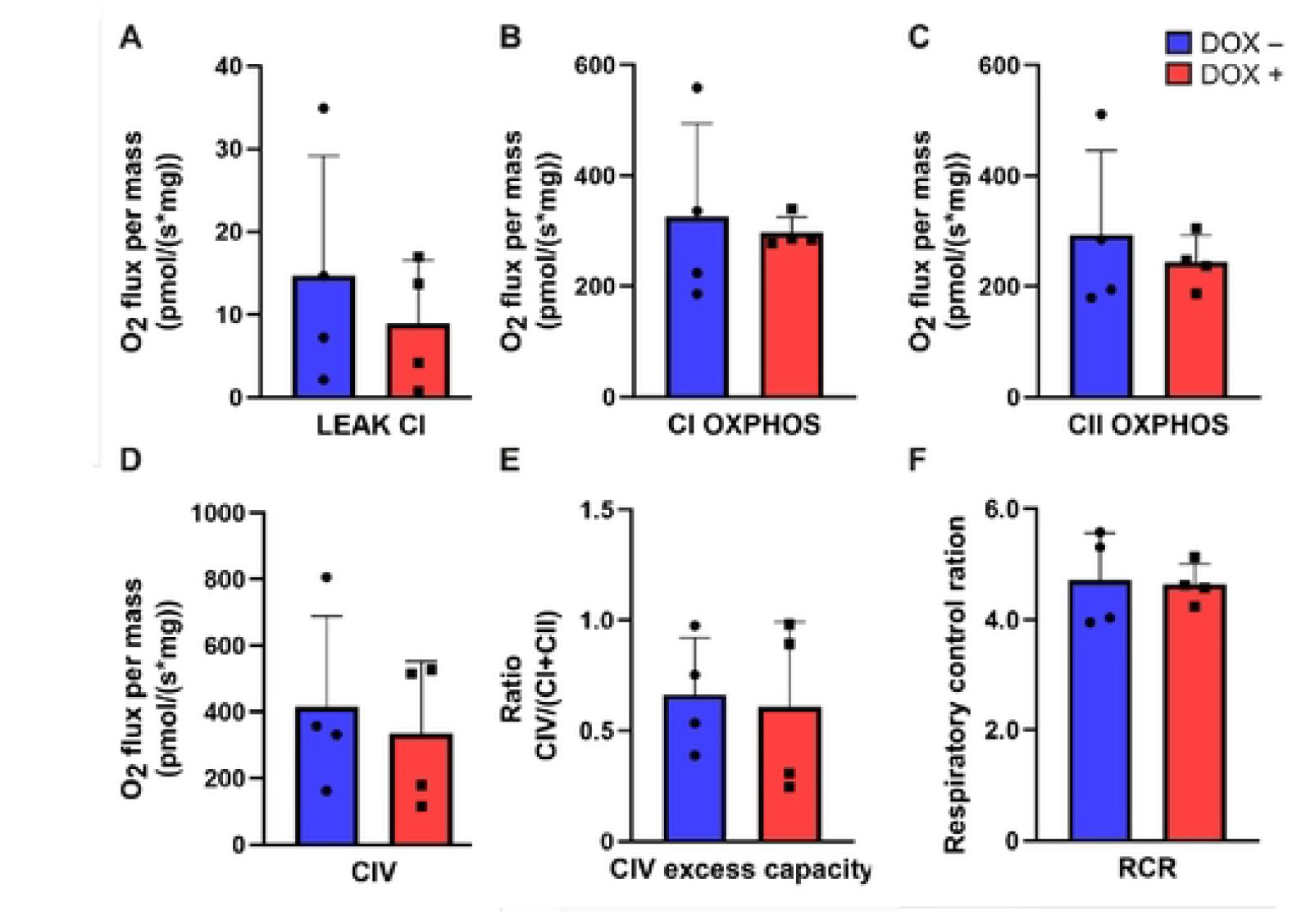
High-resolution respirometry analysis of mitochondrial function in freshly isolated soleus muscle of DOX– and DOX+ muscle-specific Cbr4 KO mice using the Oroboros Oxygraph-2k. Mitochondrial respiration was assessed by sequential addition of substrates and inhibitors. Complex I (CI) linked LEAK respiration was measured with glutamate and malate, followed by ADP to determine CI-supported OXPHOS. CI was then inhibited with rotenone, and succinate was added to assess complex II (CII) supported OXPHOS. Complex III was inhibited with antimycin A, after which complex IV (CIV) respiration was stimulated using TMPD/ascorbate. Outer mitochondrial membrane integrity was verified by cytochrome c addition. (A) Complex I (CI) linked LEAK respiration. (B) CI supported oxidative phosphorylation (OXPHOS) capacity. (C) Complex II (CII) supported OXPHOS capacity. (D) Complex IV (CIV) supported respiration. (E) CIV excess capacity relative to maximal electron transport system activity. (F) Respiratory control ratio (RCR), calculated as the ratio of OXPHOS to LEAK respiration. The data were analyzed using the unpaired t tests with Welch’s correction. Data are expressed as mean (±) SD and normalized to wet gram of soleus muscle. *n* = 3 for DOX- and n= 4 for DOX+.

### KO of *Cbr4* KO in muscle does not interfere with the lipoylation process

Lipoic acid generated by mtFAS is used as a cofactor for the lipoylation of the lipoic acid-dependent enzymes in mitochondria. In previous studies, lipoic acid polyclonal antibody that recognized the protein-bound lipoic acid has been successfully used to detect lipoylation level of E2 subunit of PDH (DLAT-LA) and KGDH (DLST-LA) (3,9). Therefore, this antibody was used to detect lipoylation of E2 subunit of PDH and KGDH in the quadriceps muscle of the DOX– and DOX+ mice. The results showed that there were no defects in the lipoylation process of the lipoic acid-dependent proteins (Fig. 5A) indicating that a *Cbr4* KO did not interfere with the generation of octanoic acid required for the lipoylation process of the mtFAS in mouse quadriceps muscle.

**Fig. 5.**
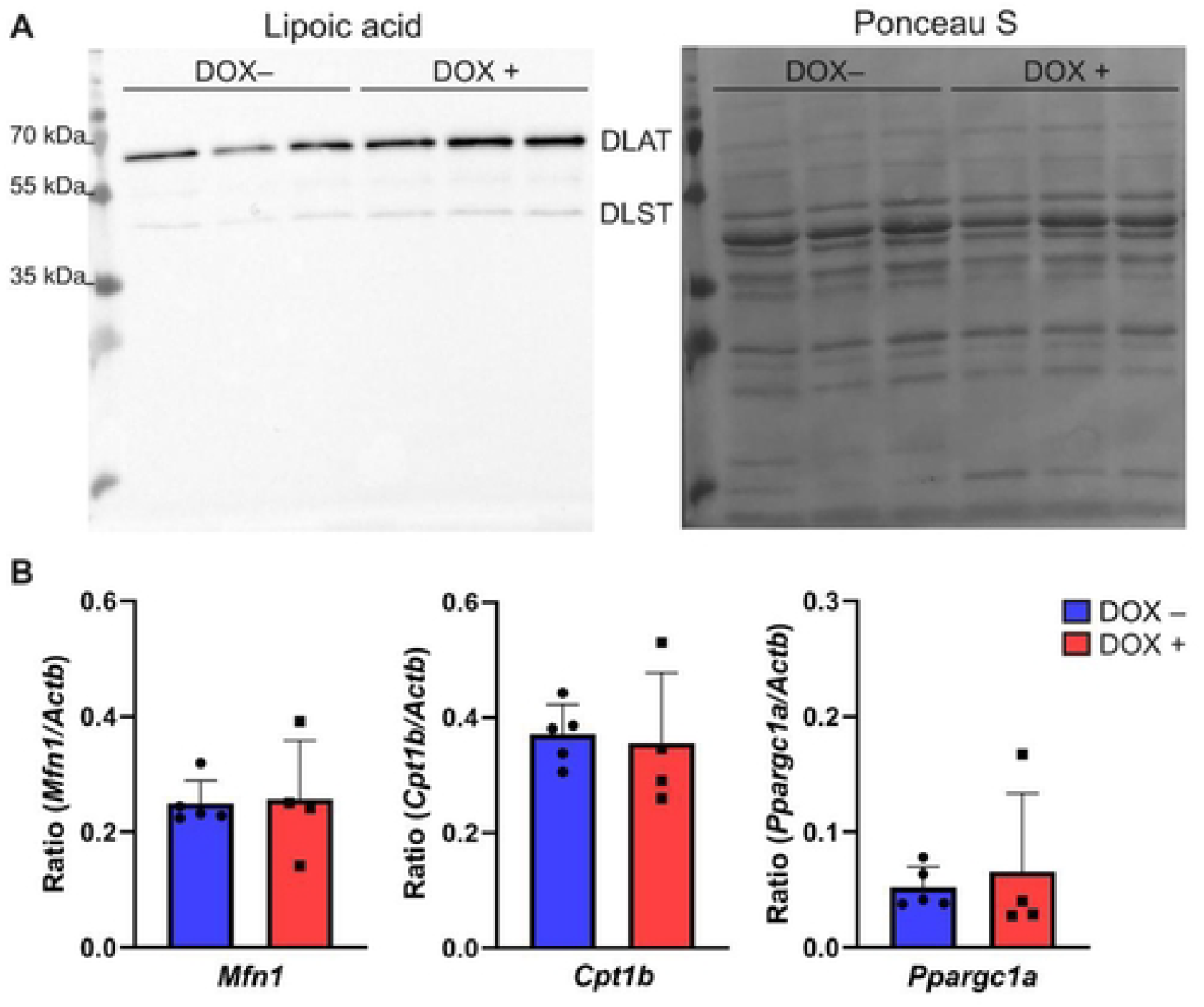
Molecular effects of muscle-specific *Cbr4* KO. (A) Immunoblot analysis for lipoylated protein in the quadriceps muscle. Lipoylated DLAT and DLST are the E2 subunits of PDH and KGDH, respectively. Immunoblotting showed that lipoylation was not defective following *Cbr4* KO. Ponceau S staining was used as a loading control. (B) mRNA expression analysis of *Cpt1b, Mfn1* and *Ppargc1α* in the quadriceps muscle of DOX– and DOX+ mice. Transcript levels were normalized against the actin (*Actb*) and presented as target gene/*Actb* ratio. No changes in mRNA expression were found for these genes in DOX+ *Cbr4* KO mice (n=4) compared to DOX– controls (n=5). Groups were compared using a Wilcoxon signed-rank test, with p< 0.05 considered as significant. Data are presented as mean (±) SD.

### Gene expression level of *Mfn1, Ppargc1a* and *Cpt1b* in Cbr4 KO muscle

Mitochondria are dynamic organelles with the ability to divide or fuse. Mitofusin (MFN1 and MFN2) proteins coordinate the fusion of the outer mitochondrial membrane (OMM) (12). Morphological differences in mitochondria have been reported for both depletion and overexpression of several mtFAS components (13–15). To determine if fusion might be affected by the *Cbr4* KO, *Mfn1* expression levels were measured in the DOX– and DOX+ mice. However, ddPCR analysis did not show differences in the *Mfn1* expression level between the control and *Cbr4* KO samples (Fig. 5B). Deletion of mitochondrial β-oxidation components in mice was previously found to be involved in upregulation of genes involved in mtFAS pathway. One of the identified genes was *Cbr4* (16). Additionally, this study found increased expression of *Ppargc1α*, which has been liked to a muscle fiber switch (16–18). Therefore, we tested transcription levels of *Ppargc1α* and *Cpt1b*, which encodes an enzyme of the long-chain β-oxidation in muscle mitochondria (19). Our ddPCR data showed no difference in *Cpt1b* and *Ppargc1α* gene expression between the DOX– and DOX+ quadriceps muscle samples (Fig. 5B).

## Discussion

Here, we have shown that CBR4 is essential for embryonic development in mice. This is consistent with previous results indicating that the mtFAS pathway is essential for mammalian embryonic development [10]. Complete ablation of MECR, resulting from the deletion of exon 2 of the Mecr gene, causes mouse embryos to perish prenatally between E9.5 and E 11.5. Embryonic lethality studies of these mice showed abnormal placental development and cardiac abnormalities (9). In another mouse mtFAS dysfunction model, mice homozygous for loss of MCAT (Malonyl CoA-acyl carrier protein transacylase) were not viable (21). All in all, our result that breeding *Cbr4*^*tm1d*^ mice did not produce homozygous KO pups is consistent with these previous mtFAS dysfunction mouse model studies, indicating the involvement of CBR4 in early mammalian development. As CBR4 appears to be of similar importance to mitochondrial function as other mtFAS components, the emergence of patients with CBR4 dysfunction and symptoms resembling MEPAN syndrome can be expected.

In contrast, the skeletal muscle-specific *Cbr4* deletion showed no discernable effect on the parameters we investigated. Although our experiments started with nine *Cbr4* KO candidates, we only identified four (female) KO mice with confidence. Lacking a specific CBR4 antibody, our toolbox was restricted to the investigation of Cbr4 transcript levels. Our inability to confirm the detection of males carrying the deletion mutations with any confidence using ddPCR may be attributed to already very low levels of *Cbr4* expression in male mice, leading to difficulties in detection of meaningful differences and hence an insurmountable artifact of our gene expression analysis in males. We cannot completely exclude the possibility of inefficient activation of the *Cre* driver construct in several (male) mice, although we find this scenario unlikely. The muscle-specific gene KO approach we used is well documented in previous work (11). Although in several more of the mice exposed to DOX, the KO event may have successfully occurred, we chose to err on the side of caution and to only analyze the group of female mice for which we were certain that *Cbr4* was inactivated in the skeletal muscle.

In contrast to the complete KO, skeletal muscle-specific *Cbr4* KO mice did not lead to any detectable defects in physical performance. Overall growth, feeding, weight gain and grip strength were indistinguishable between DOX– and DOX+ mice (Fig. 2A and 2B). The available literature showed that tissue-specific knockout/knock-in of the mtFAS component causes variable phenotypes depending on the target tissue. Purkinje-cell (PC)-specific *Mecr* KO mice causes loss of PCs due to absence of MECR protein in these cells, resulting in a neurodegenerative phenotype. PCs from these cells display mitochondrial morphological abnormalities, hypolipoylated target proteins as well as reduced mitochondrial OXPHOS. (20). Mice carrying a tamoxifen-inducible, ubiquitously targeted conditional *Mcat* construct exhibit severely reduced body weight, develop alopecia and kyphosis and have reduced grip strength and stamina upon tamoxifen induction of the KO. However, motor coordination and balance of these KO mice remain intact. Interestingly, no morphological changes in the muscle, bone, or connective tissue in the spinal column or hind limps were observed. The lipoylation level and enzymatic activities of the lipoic acid-dependent enzymes, however, were markedly reduced in the tissues studied. In contrast to the ubiquitous *Mcat* KO, the heart muscle-specific *Mcat* KO mice did not show any phenotype although protein lipoylation in the heart muscle was compromised but not abolished (21). Interestingly, and in line with our results, no direct effects on skeletal muscle strength or heart function have been reported MEPAN or MCAT deficient patients (3,22,23).

Knockdown of mitochondrial ACP in cultured HEK cells causes loss of lipoylation which could not be rescued by supplementing the media with exogenous lipoic acid (24), and lipoic acid feeding to PC specific KO *Mecr* mice could not prevent the development of neurodegeneration due to mtFAS defect (25), indicating a requirement for that endogenous lipoic acid synthesis in mammals. Our immunoblot analysis of DOX– and DOX+ skeletal muscle-specific *Cbr4* KO mice showed no differences in the level of DLAT-LA and DLST-LA (Fig. 5A). This is, however, in accord with observations on extracts obtained from cultured fibroblasts from MEPAN patients, where lipoylated proteins could be detected although the MECR protein was absent (3).

In our skeletal muscle-specific *Cbr4* KO mouse model we did not observe any mitochondrial respiration defects (Fig.4). Deletion of components of mtFAS in budding yeast causes loss of respiratory ability, evident from the inability to grow on non-fermentable carbon source (1). Reduction of the maximum capacity of electron transport system (ETS) was reported for fibroblasts of some MEPAN patients (3). However, fibroblasts from a MEPAN patient with strongly reduced MECR protein showed no difference in abundance of OXPHOS complexes and only a modest decrease of CI and CIV enzymatic activities (3). The analysis of the mitochondrial respiration in the tissues of mice subjected to a post-partum ubiquitous *Mcat* KO showed that the effects of disrupted mtFAS on the respiration capacity were not universal. There was no defect in the respiration in the kidney mitochondria, and in skeletal muscle only late-stage mild defects could be observed in the ETC (21).

Patients with *MECR* pathogenic variants suffer predominantly from neurological defects. Their cognitive skills, cardiac and muscle functions usually remain intact (3). Deletion of the mitochondrial β-oxidation very long-chain acyl-CoA dehydrogenase (VLCAD) enzyme from skeletal muscle of mice induces a metabolic adaptation to glycolysis in response to energy demand in mice (16). qPCR analysis of quadriceps muscle showed concomitant upregulation of the *Cbr4* gene along with other genes involved in the mtFAS pathways. Additionally, this study found increased expression of *Ppargc1α*, which has been correlated with a switch from type II to type I muscle fibers and to the proliferation and activation of oxidative metabolism (16–18,26). Conversely, the pathogenic mutation in the mtFAS gene *ACSF3* causes upregulation of mitochondrial β-oxidation genes (27). Therefore, we used our skeletal muscle-specific *Cbr4* KO mouse model to see if *Cpt1b*, encoding the rate limiting enzyme for long-chain β-oxidation in muscle (19,28), and *Ppargc1α* gene expression is altered in DOX+ mice. We did not detect changes in the expression levels of these genes (Fig. 5B).

Our data on the function of CBR4 in mice confirms that the mtFAS pathway is indispensable in the mammalian developmental process and dispensable in a tissue specific context. MEPAN patient symptoms appear to be restricted to neurodegeneration effects. At first sight, the absence of any effect of a muscle-specific disruption of mtFAS on the function of this tissue/organ is puzzling. However, our results match those of previous studies on *Mcat* inactivation in mice (21) and data obtained with patient cells (3). We have previously hypothesized that the neuron-specific phenotype of mtFAS disorders may be due to the absence of mitochondrial β-oxidation in neurons (29). According to this idea, tissues that are capable of fatty acid breakdown in mitochondria may be able to use mitochondrial β-oxidation intermediates as an alternative source of attachment of acyl groups to ACP, reducing the requirement for functional mtFAS. Our results of the skeletal muscle-specific *Cbr4* KO are consistent with such a hypothesis.

## Materials and Methods

### Study approval and mouse strains

Mice were maintained in the experimental animal center of University of Oulu, Oulu, Finland according to accepted criteria for humane care and use of experimental animals. Mice were housed as a group in standard cages at 55% humidity, 22°C temperature and a 12 h/12 h dark/light cycle (lights on from 7:00 a.m. to 7:00 p.m.). Food and water were provided *ad libitum* following the guidance of European Union directive 2010/63/EU for animal wellbeing. The experimental protocols were approved by the Animal Care and Use Committee of National Animal Experiment Board, Finland (license number ESAVI/9204/2020).

### Generation of *Cbr4* knockout mice

“Knockout-first” type *Cbr4* mutant mice were generated from embryonic stem (ES) cells obtained from European Conditional Mouse Mutagenesis Program (EUCOMM). Briefly, floxed *Cbr4* mice were generated by standard techniques using a targeting vector containing a neomycin (G418) resistance cassette flanked by *FRT* sites. Exon 2 of the *Cbr4* gene was inserted into two flanking *loxP* sites. After electroporation of the targeting vector into ES cells (C57BL/6N-A/a), G418-resistant ES cells were screened with PCR with *Cbr4* gene specific primer targeting exon 2 and primer adjacent to the downstream *loxP* site to ensure the clone has intended allele and the 3’ *loxP* site is present. The recombinant ES cells were microinjected into blastocysts from C57BL/6JOlaHsd. Two chimeric pups were born based on PCR-analysis and coat color. Mice were crossed with C57BL/6JOlaHsd / C57BL/6NCrl background. *FRT*-flanked selection cassette between exons 1 and 2 was deleted by breeding the mice with transgenic mice Tg(*ACTFLPe*)9205Dym/J, expressing the *FLPe* recombinase gene under direction of the human *ACTB* promoter (*ACTB::FLPe*) (30). The resulting homozygous floxed *Cbr4* (*Cbr4*^tm1c^) mice carried a conditional knockout cassette with *Cbr4* exon 2 flanked by *loxP* sites. The *Cbr4*^tm1c^ strain mice were bred with Tg(*CAG-Cre*)13Miya mice with ubiquitous *Cre* expression under the control of cytomegalovirus immediate early enhancer-chicken β-actin hybrid (CAG) promoter (31) to generate *Cbr4*^tm1d^ mice in which the floxed exon 2 was deleted.

To obtain inducible skeletal muscle-specific KO mouse line, *Cbr4*^tm1c^ mice were bred with B6;C3-Tg (ACTA1-rtTA, tet0-cre)102Monk/J (also knowns as HSA-rtTA/TRE-Cre) mice from the Jackson Laboratory (11). Pups were fed with 2 mg/ml of a tetracycline analog DOX supplemented with 5% sucrose starting at the age of 6 weeks and continuing for 3 weeks. The control littermate was fed with 5% sucrose. Among the 9 male and 10 female pups, 4 of the male pups and 5 of the female pups were given 2 mg/ml DOX supplemented with 5% sucrose. The growth of the mice was monitored by weighing them twice in a month over a 12-month period.

### Grid hanging test

The grid hanging test was done following a protocol similar to a recently published article (25). To perform the grid hanging test, a metal wire grid with a diameter of 1×1 cm was used. The mouse was placed in the center of the grid. Then the grid was inverted, and a stopwatch was used to record the time. The elapsed time was recorded either when mouse dropped from the grid on a soft bedding or at the end of five-minute time and the test was repeated three times. This test was done twice in a month over a 12-month period.

### Grip strength analysis

The force of forepaws was analyzed with Grip Strength Meter (Columbus instruments). First, mice grasp the bar of the instrument and then the animal is gently pulled away from the device. The maximal force before mice releases the bar is detected. The experiment was repeated three times, and the biggest force was used for the analysis.

### Running tests

Prior to running test on treadmill (Panlab treadmill, Harvard apparatus) the mice were accommodated the treadmill and running three times. The running test for 14 minutes was initiated with speed of 8 cm/s for 2 minutes, then 2 minutes on speed 10 cm/s and finally 10 minutes on speed 12 cm/s. Altogether the running distance was 93 meters. Blood glucose and lactate concentrations were monitored from the *vena saphena* using a glucometer (mylife Pura, SteriLance Medical) and a lactometer (Lactate Scout +, SensLab/EKF Diagnostics). Blood glucose and lactate levels were analyzed before, right after and 30 minutes after the running exercise. Two weeks later another running test was performed on the same treadmill. Here the test was initiated with speed of 8 cm/s for 2 minutes. After this the speed was increased by 3 cm/s every second minute until maximum speed of 50 cm/s. The exercise continued until the mice stopped running consecutively.

### Skeletal muscle collection

At the end of the behavioral analysis, mice were euthanized by exposure to CO_2_ followed by a cervical dislocation. Quadriceps and soleus muscles were isolated from DOX– and DOX+ mice. For RNA isolation, the quadriceps muscle was immediately separated from the euthanized mice, transferred to RNAlater stabilization reagent (Qiagen) and flash frozen in liquid nitrogen. For western blot, the collected quadriceps muscle was flash frozen in liquid nitrogen. The samples were stored in –70°C.

### Droplet digital PCR (ddPCR) to detect gene expression

ddPCR experiment was performed to detect the gene expression level in the skeletal muscle-specific *Cbr4* KO mice. Total RNA from quadriceps muscle was collected with a RNeasy Mini Kit (Qiagen) according to the manufacturer protocol. In-column DNase treatment done for all samples using RNase-Free DNase Set (Qiagen). cDNA from total RNA was generated with a RevertAid First Strand cDNA Synthesis Kit (ThermoFisher Scientific) using the random hexamer primer (supplied with the kit) following manufacturer protocol. The synthesized cDNAs were used as a template for the ddPCR experiment using ddPCR gene expression probe assay. Unless otherwise stated, all the instruments, software, consumables, primers, and probes related to the ddPCR experiments were purchased from Bio-Rad. The gene specific primers used in the study are listed in (Table 1). The mouse actin gene was used as an endogenous control for the normalization of gene expression. The actin primers and probe sequence were purchased from the ddPCR™ gene expression assay (assay ID dMmuCPE5195285). Each of the ddPCR reaction mixtures consists of 22 μl of master mixture with 10 μl of 2x ddPCR Supermix for probes (no dUTP). Droplet emulsions were generated using the QX200 droplet generator. 40 μl of emulsions were carefully transferred to the 96-well plate and heat-sealed with pierceable foil using the PX1 PCR plate sealer. The sealed 96-well plate transferred to the T100™ Thermal Cycler. The thermocycling condition was as follows, 95°C for 10 minutes followed by 40 cycles at 94°C for 30 seconds, 55°C for 1 minute. The enzyme deactivation step was done at 98°C for 10 minutes. The ramp rate was 2°C/second. After thermocycling, the 96-well plate was kept at 4°C overnight. QX 200TM droplet reader was used to count the droplets utilizing the HEX/FAM channel. QuantaSoft 1.7.4 software was used to analyze gene expression levels, and this is expressed as ratio of target gene/actin. Three technical replicates were used in the analysis.

### Western blot analysis of the quadriceps muscle

Total proteins were extracted from quadriceps muscle of the DOX– and DOX+ mice using RIPA lysis buffer (1M NaCl, 1% Igepal ca-630, 0.5% sodium deoxycholate,0.1% SDS, 50mM Tris-Cl pH 7.4). Protease inhibitor tablet (cOmplete, EDTA free, Roche) was added in the RIPA buffer immediately before lysis. Protein concentration was measured using the Bio-Rad protein assay (Bio-Rad). SDS-PAGE electrophoresis was done using the 12% SDS-PAGE gel. Each well of the gel was loaded with approximately 40 μg of protein samples. After the gel run, the gel was blotted on a 0.2 μm nitrocellulose membrane using Bio-Rad Trans-Blot Turbo system and Trans-Blot Transfer Pack. Efficient protein transfer to the membrane was tested immediately after transfer using the Ponceau S staining and imaged was taken as a loading control. The membrane was washed with TBST buffer (50 mM Tris, 150 mM NaCl, 0.05% Tween 20) and blocked overnight at 4^°^ C in 1× Casein Blocking Buffer (Sigma-Aldrich). After washing the membrane with TBST buffer, western blotting was done with anti-lipoic acid antibody (1:2500, EMD Millipore) for 1 hour at room temperature. HRP-conjugated goat anti-rabbit IgG (Bio-Rad) antibody at a dilution of 1:5000 was used as a secondary antibody. The signal was detected from the membrane using Clarity ECL reagent (Bio-Rad) with a 5-minutes incubation time. The membrane was imaged using Bio-Rad XRS camera.

### Histological analysis of the quadriceps muscle

Quadriceps muscles were fixed in 4% paraformaldehyde in 100 mM NaPi buffer (pH 7.4) over night and embedded in parafilm. For hematoxylin and eosin (H&E) analysis, 5 μm section of both DOX– and DOX+ mice were used. The stained sections were scanned using the NanoZoomaer S60 slide scanner at 20X mode in the transgenic and tissue phenotyping core facility of University of Oulu, Finland. For fiber cross sectional area (FCSA) measurement, the transverse section of each slide was used. For each section, three areas were selected for quantification. Cellpose-SAM (32) was used for segmentation of individual fibers and the fiber area was quantified using ImageJ (v1.54). The average area per fiber of the three selected regions was used as representation of the section.

### Mitochondrial respiratory complex activity studies

Mitochondrial respiratory complexes activities were measured using the high-resolution respirometry Oxygraph-2K (OROBOROS) in soleus muscle collected from DOX– and DOX+ mice. The OROBOROS instrument contains two identical glass chambers with cathodes. Each chamber has two milliliter (ml) respiratory media with continuous stirring at 37°C. The complex activity measurement was performed following a previously published protocol (33). Soleus muscles were collected from the euthanized mice and placed on the petri dish containing ice-cold isolation solution A for permeabilization of muscle fibers (10 mM Ca-EGTA buffer (free concentration of calcium 0.1μM) 20 mM imidazole, 20 mM taurine, 49 mM K-MES, 3 mM KP_i_, 9.5 mM MgCl_2_, 5.7 mM ATP, 15 mM phosphocreatine, 1 μM leupeptin pH 7.1). For the permeabilization of the muscle tissue saponin at a concentration of 50 μM in 2 ml permeabilization solutions A was used and mixed gently at 4°C for 20 minutes. After that permeabilized fibers were washed 3 times at 4°C for 5 minutes in mitochondrial respiratory medium (0.5 mM EGTA, 3 mM MgCl_2_, 20 mM taurine, 10 mM KP_i_, 20 mM HEPES, 1 g liter-1 BSA, 60 mM potassium-lactobionate, 0.3 mM dithiothreitol, pH 7.1) to remove the saponin, ADP, and ATP.

After equilibrating the Oxygraph chamber with 2 ml of mitochondrial respiratory medium buffer, the permeabilized fibers were transferred to the chamber and chambers were closed. Resting CI supported respiration was measured by adding 10 mM glutamate and 5 mM malate. Then, ADP at a final concentration of 2 mM was added to get state 3 maximum mitochondrial respiration. Followed by addition of 0.5 μM of rotenone for the inhibition of CI activity. After that, 10 mM succinate was added to the chamber for the induction of CII respiration. This step is followed by the inhibition of CIII by addition of 5 μM antimycin A. In the next step, 0.5 mM TMPD and 2 mM ascorbate were used to measure CIV activity. To check the OMM intactness and as a quality control for the permeabilized preparation, 10 μM cyt C was added to the chamber. Respiration rates were normalized using dry weight of the muscle fibers and oxygen consumption rate is expressed per mg of dry muscle weight. Datalab software V 4.3.4.70 from OROBOROS instruments were used for the data analysis and statistical analysis was done using the GraphPad prism 9 software.

## Author contributions

J. Kalervo Hiltunen (J.K.H.), and Alexander J. Kastaniotis (A.J.K.) conceptualized and designed the study. A.J.K, Kaija J. Autio (K.J. A), and Guangyu Jiang (G.J.) generated and analyzed the knockout mice. Ali J Masud (AJM) participated in the mouse tissue collection, performed ddPCR analysis, western blot analysis and tissue staining. AJM and M. Tanvir Rahman (MTR) carried out the OROBOROS Oxygraph-2K experiment, and data analysis. Irene M.G.M. Hemel (IH) performed data analysis for ddPCR, western blot and tissue staining. AJM, IH and KJA generated all the figures and tables. AJM wrote the draft manuscript. JKH, AJK, KJA, GJ, MTR and IH contributed to manuscript revision. J.K.H. and A.J.K. supervised the study.

## Acknowledgements

This work was made possible by the support from the Jane and Aatos Erkko Foundation (Finland), The Academy of Finland (Now “Research Council Finland”) and the Sigrid Jusélius Foundation (Finland). The research was carried out with the support of Biocenter Oulu, Transgenic and Tissue Phenotyping Core Facility, endowed by the University of Oulu, Finland and Biocenter Finland and the Oulu Laboratory Animal Centre Research Infrastructure, University of Oulu, Finland.

## Notes

### Competing Interest Statement

The authors have declared no competing interest.

